# Establishing the contribution of active histone methylation marks to the aging transcriptional landscape of Drosophila photoreceptors

**DOI:** 10.1101/2022.09.30.510348

**Authors:** Juan Jauregui-Lozano, Kimaya M. Bakhle, Arrianna C. Hagins, Vikki M. Weake

## Abstract

Studies in multiple organisms have shown that aging is accompanied by several molecular phenotypes that include dysregulation of chromatin. Since chromatin regulates DNA-based processes such as transcription, alterations in chromatin modifications could impact the transcriptome and function of aging cells. In flies, as in mammals, the aging eye undergoes changes in gene expression that correlate with declining visual function and increased risk of retinal degeneration. However, the causes of these transcriptome changes are poorly understood. Here, we profiled chromatin marks associated with active transcription in the aging *Drosophila* eye to understand how chromatin modulates transcriptional outputs. We found that both H3K4me3 and H3K36me3 globally decrease across all actively expressed genes with age. However, we found no correlation with changes in differential gene expression. Downregulation of the H3K36me3 methyltransferase Set2 in young photoreceptors revealed significant changes in splicing events that overlapped significantly with those observed in aging photoreceptors. These overlapping splicing events impacted multiple genes involved in phototransduction and neuronal function. Since proper splicing is essential for visual behavior, and because aging *Drosophila* undergo a decrease in visual function, our data suggest that H3K36me3 plays a role in maintaining visual function in the aging eye through regulating alternative splicing.

## INTRODUCTION

Eukaryotic genomes are organized into a nucleoprotein structure termed chromatin that governs access to the underlying DNA, and thereby plays a key role in regulating processes such as transcription, replication, and DNA repair ^1,2^. During aging, chromatin structure often becomes dysregulated, and these changes are considered to be one of the nine “Hallmarks of Aging” ^3^. In the aging eye, cells in the retina undergo changes in DNA methylation and gene expression that correlate with decreased visual function and increased cell death ^3–7^. Moreover, epigenetic mechanisms are implicated in the development and pathology of ocular diseases such as age-related macular degeneration ^8,9^. These data suggest that the disruption of the chromatin landscape in the aging eye contributes to changes in its transcriptome, potentially increasing the risk of ocular disease.

Histone methylation is one of the most prevalent and well characterized chromatin modifications, and plays an important role in regulating processes including transcription and splicing ^10,11^. During different stages of the transcription cycle, the N-terminal tail of histone H3 is subject to tri-methylation on two specific lysine residues, Lys-4 (H3K4me3) and Lys-36 (H3K36me3) ^12–14^. Thus, H3K4me3 and H3K36me3 mark actively expressed genes and their levels broadly correlate with transcript levels in many cell types including photoreceptors ^15,16^. *Drosophila melanogaster* provides a simplified model in which to study the role of these active histone methyl marks because there is less redundancy in the enzymes that deposit these marks relative to humans. In flies, there are four enzymes that methylate H3K4: Ash1 generates H3K4me2, while Trx, Trr (human MLL3/4), and Set1 (human SET1A/B) are responsible for tri-methylation of H3K4 ^17^, with Set1 depositing the bulk of global H3K4me3 in flies ^18,19^. In contrast, only two enzymes methylate H3K36: NSD is responsible for H3K36me2, while Set2 adds the third methyl group to generate H3K36me3 ^17,20,21^.

Previous microarray based chromatin-immunoprecipitation (ChIP-CHIP) analysis in *Drosophila* heads suggested that H3K4me3 and H3K36me3 levels decreased with age ^22^; in contrast, no decreases in repressive histone methyl marks such as H3K27me3 and H3K9me3 were reported ^22,23^. H3K4me3 and H3K36me3 have been linked to the regulation of transcription initiation and elongation, respectively. Since photoreceptors experience age-associated changes in transcription factor activity ^24,25^, as well as splicing ^26,27^, we sought to study how histone methylation contributed to the aging transcriptome. Further, we wondered if decreasing levels of one of these active marks would directly contribute to any of the age-associated changes observed in the transcriptome of aging photoreceptors. Our study reveals that aging photoreceptors undergo decreases in levels of both H3K4me3 and H3K36me3 at all expressed genes, which correlate with decreased chromatin accessibility at promoters, but not with age-associated changes in differential gene expression. Using photoreceptor-specific knockdown of the H3K36me3-methyltransferase, Set2, we show that decreased H3K36me3 does not contribute to age-associated changes in differential gene expression or chromatin accessibility, but instead leads to changes in alternative splicing that overlap significantly with those observed during aging and include many gene targets involved in visual function. These data demonstrate that the dysregulation of specific chromatin marks during aging can contribute to distinct effects on gene expression profiles. Thus, we propose that alternations in multiple chromatin marks in the aging eye likely disrupt cellular and tissue homeostasis and may contribute to the increases risk of ocular disease with age.

## RESULTS

### Aging photoreceptors experience genome-wide changes in H3K4me3 and H3K36me3 distribution

We sought to examine photoreceptor-specific changes in two active histone methylation marks during aging: H3K4me3 and H3K36me3. In *Drosophila*, as well as other animals, H3K4me3 is deposited around the transcription start site (TSS) of actively expressed genes (Fig. 1A), corresponding with the association of Set1-containing histone methylation complexes with the initiating RNA polymerase II. In contrast, H3K36me3, which is deposited by Set2 associated with the elongating RNA polymerase II, is enriched over the gene bodies of actively expressed genes, with higher levels towards the 3’ end of the gene (Fig. 1A). To profile these histone marks in photoreceptors, we purified photoreceptor nuclei from *Rh1-Gal4>UAS-GFP^KASH^* flies, which express Green Fluorescent Protein (GFP) fused to the nuclear membrane-anchored KASH domain (GFP^KASH^) in outer photoreceptors, as described previously ^15^. Briefly, GFP^KASH^-tagged nuclei are isolated using anti-GFP antibodies and magnetic beads, allowing immunoenrichment of nuclei from specific cell types (Fig. 1B). We previously showed that this nuclei immuno-enrichment (NIE) approach can be used with chromatin-immunoprecipitation coupled with high-throughput sequencing (ChIP-seq) to obtain high quality data ^15^. We examined H3K4me3 and H3K36me3 in photoreceptors at day 10 (young) and day 40 (old) post-eclosion because flies show changes in visual behavior and photoreceptor-specific gene expression between these ages ^4^. In addition, we had available transcriptome and chromatin accessibility data for these same time-points in this genotype, allowing us to integrate these data with the ChIP-seq analysis in the current study.

**Figure 1.**
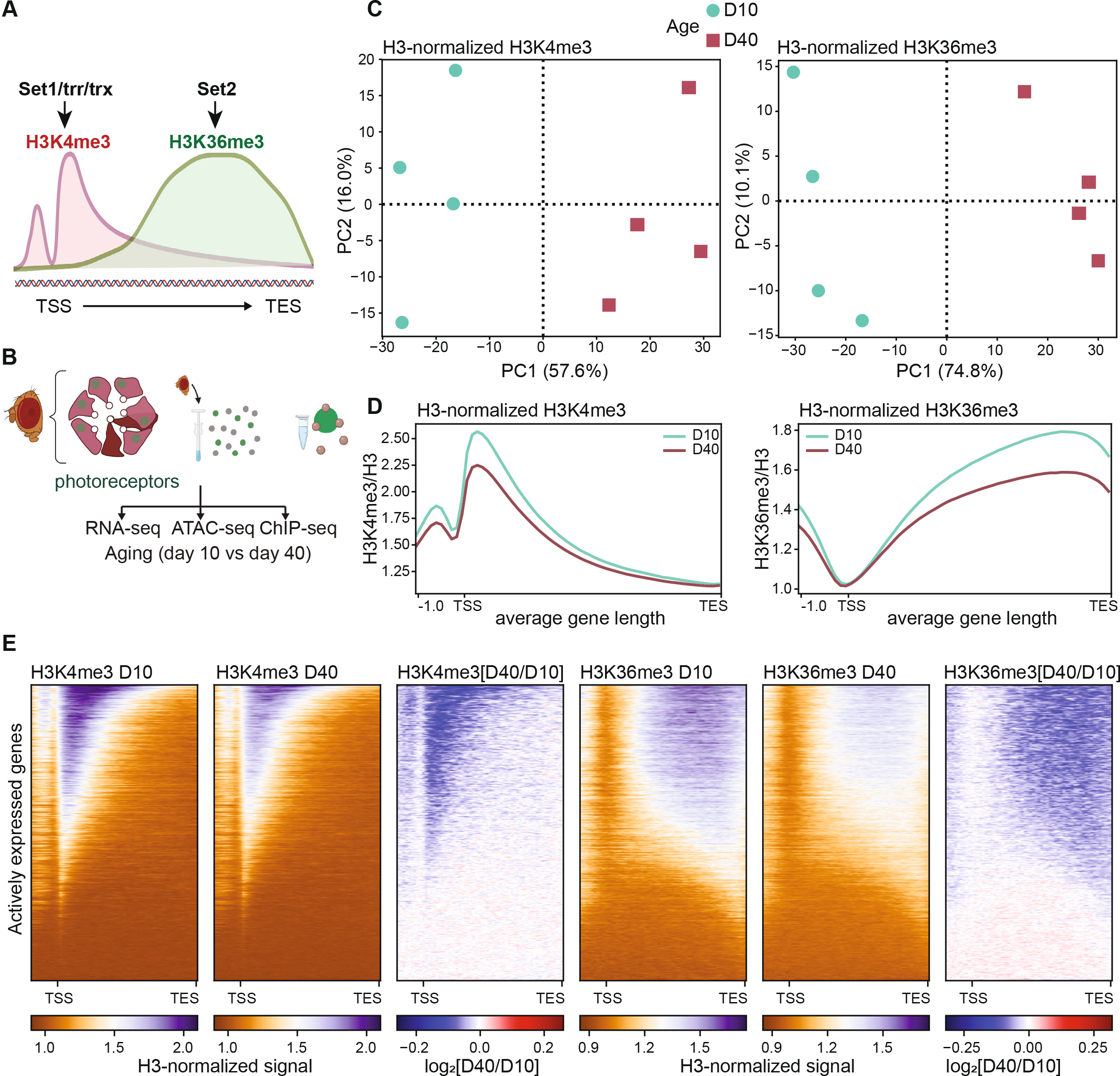
The active histone methylation landscape changes with age in photoreceptor neurons. **A**. Diagram of the spatial distribution of active histone methylations around the gene body. In *Drosophila*, H3K4me3 is deposited by three enzymes (Set1, trr, trx), while H3K36me3 is deposited by Set2. TSS, transcription start site; TES, transcription end site. **B**. Experimental diagram of nuclei immuno-enrichment protocol to purify aging photoreceptor nuclei. ChIP-seq analysis from the current study was integrated with RNA-seq and ATAC-seq from aging photoreceptors described in previously published studies. **C**. Principal component analysis (PCA) H3- and CPM-normalized H3K4me3 (left) and H3K36me3 (right) (n = 4; D10, day 10; D40, day 40). **D**. Gene metaplot of the average H3- and CPM-normalized H3K4me3 (left) and H3K36me3 (right) signal over gene bodies for young and old flies (n = 4). **E**. Heatmaps showing H3- and CPM-normalized signal for H3K4me3 or H3K36me3 at D10 and D40 at all actively expressed genes (blue-orange heatmaps, panels 1 – 2 and 4 - 5). Ratios (log_2_) of H3K4me3 or H3K36me3 changes between D10 and D40 are shown in the blue-red heatmaps in panels 3 and 6, representing the fold change signal in old relative to young flies with red indicating increase in active histone levels, and blue representing a decrease; no change is centered to white.

In addition to H3K4me3 and H3K36me3, we profiled histone H3 to account for age-associated changes in nucleosome occupancy, which has previously been observed in brain tissues and neural cells during aging ^28^. We then obtained H3-normalized H3K4me3 and K36me3 signals for each of the four biological replicates. The genome-wide distribution of the H3-normalized H3K4me3 and H3K36me3 samples by principal component analysis (PCA) revealed that both histone marks clustered by age (Fig. 1C), with 57.6% and 74.8% of the variation explained by age for H3K4me3 and H3K36me3, respectively. Previous studies assaying genome-wide levels of chromatin marks during aging have shown there can be global changes associated with age ^22,23,29,30^. To address this possibility, we had incorporated *Arabidopsis thaliana* chromatin to perform spike-in normalization (Supp. Fig. 1A), which can improve data analysis of experiments where global changes in signal are expected ^31,32^. However, in our hands, we observed similar age-dependent differences in genome-wide distribution of the H3-normalized H3K4me3 and H3K36me3 data whether or not we used the spike-in normalization (Supplemental Fig. 1B), suggesting that this normalization approach was unnecessary for these data. Taken together, these data show that the active histone landscape of photoreceptor neurons changes with age.

### Aging photoreceptors experience a genome-wide decrease of histone methyl marks at actively expressed genes

H3K4me3 and H3K36me3 are deposited by enzymes that associate with the actively transcribing RNA polymerase II, and are considered marks of active transcription ^12,33^. Thus, we evaluated levels of these marks over the gene bodies of actively expressed genes, based on expression levels of more than 5 transcripts per million (TPM) expression as quantified by RNA-seq ^25^. As expected from previous ChIP-seq analysis of these marks in young photoreceptors, gene metaplots showed the characteristic enrichment patterns expected for H3K4me3 around the TSS, and H3K36me3 over gene bodies, with enrichment towards the 3’ end of genes. We found an age-associated decrease in total H3-normalized H3K4me3 and H3K36me3 signal (Fig. 1D), resembling the previous decrease in H3K4me3 and H3K36me3 levels observed using ChIP-CHIP analysis of aged *Drosophila* heads ^22^. Importantly, this age-associated decrease was apparent with both merged (Fig. 1D) and individual biological replicates for each time point (Supp. Fig. 1C), and was still visible with spike-in normalization (Supp. Fig. 1D).

To test if this age-associated decrease in histone methyl marks was driven by a subset of genes, we obtained heatmaps of H3-normalized H3K4me3 and H3K36me3 signal at day 10 and day 40, where genes are sorted vertically based on their signal enrichment (Fig. 1E: orange-blue heatmaps). To quantify the loss of signal, we obtained log_2_-transformed ratios of H3K4me3 or H3K36me3 at day 40 divided by the corresponding signal at day 10 (Fig. 1E, log_2_ ratios: blue-red heatmap). These heatmaps revealed that nearly all actively expressed genes that were marked by either H3K4me3 or H3K36me3 showed a significant loss of signal with age. Based on these data, we conclude that *Drosophila* photoreceptors show a global decrease in active histone methylation marks during aging.

Since histone methylation levels are regulated by both their deposition by histone writers, and removal by histone erasers ^34,35^, we next assessed if genes involved in maintenance of H3K4me3 and H3K36me3 showed changes in gene expression during aging, as measured by RNA-seq. We did not observe any significant decrease at the transcript level for the methyltransferases that deposit H3K4me3 or H3K36me3, and only one of the demethylases involved in H3K36me3 demethylation showed an increase in expression with age (Supp. Fig. 2).

**Figure 2.**
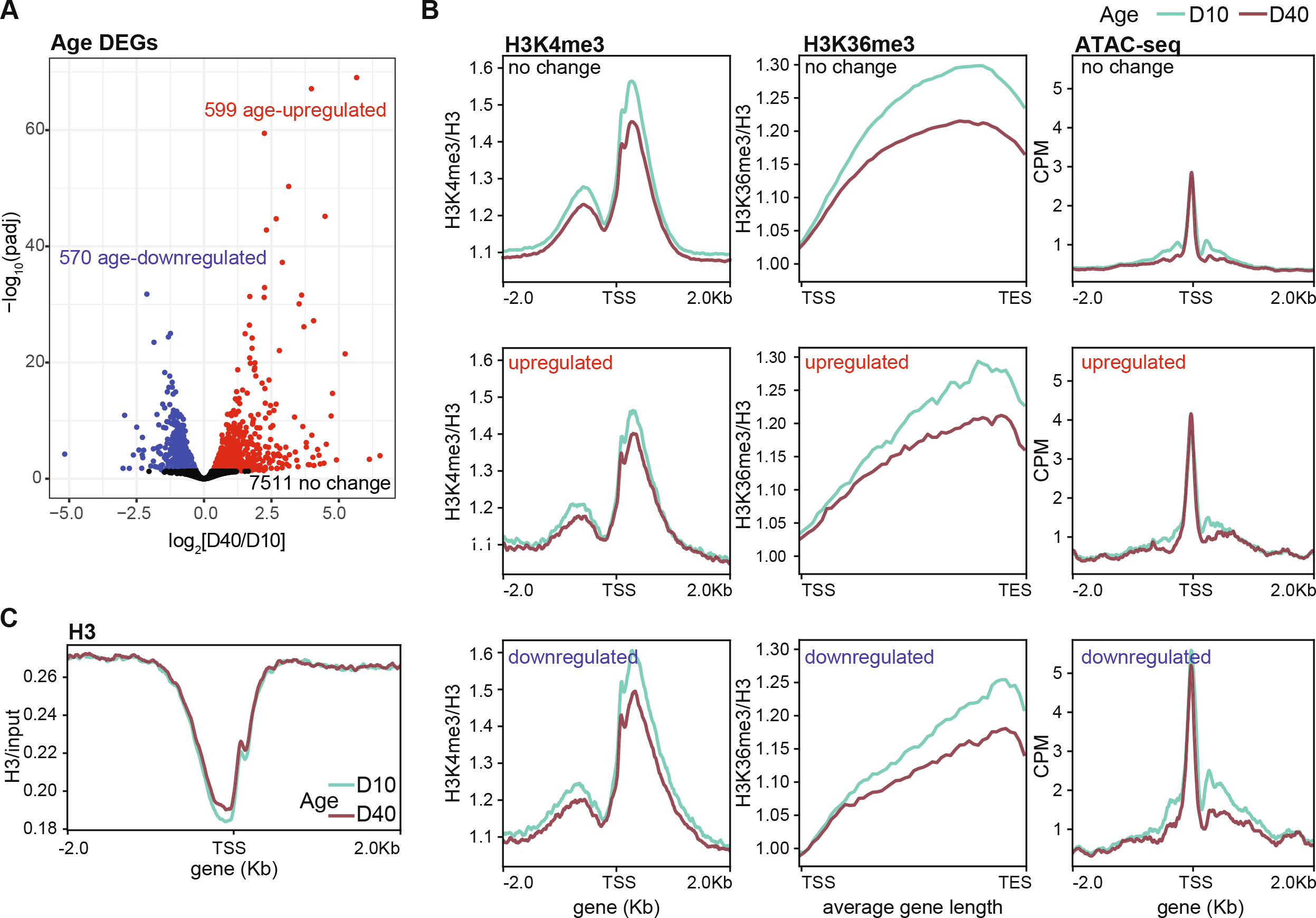
Actively expressed genes have decreased active histone methyl marks independent of their transcriptional status. **A**. Volcano plot showing genes that were differentially expressed between D10 and D40 in photoreceptors. **B**. Gene metaplots showing average H3-normalized H3K4me3 and H3K36me3 levels and ATAC-seq signal around the TSS (H3K4me3 and ATAC-seq) or over gene bodies (H3K36me3) at D10 and D40 for genes classified based on their differentially expression status with age as described in panel A. **C**. Gene metaplot of average chromatin input-normalized H3 signal (n = 4) for actively expressed genes with signal centered around the TSS.

Further, there were no significant changes in protein levels of the three enzymes detected by proteomic analysis of the aging eye: trithorax *(Trx-FBgn0003862*), Lysine demethylase 5 *(Lid-FBgn31759)*, and Lysine demethylase 2 (*Kdm2-FBgn0037659*) ^36^. These data suggest that the decreased levels of H3K4me3 or H3K36me3 in aging photoreceptors are not simply caused by altered expression of the enzymes that deposit or remove these marks.

Taken together, we conclude that *Drosophila* photoreceptors show a global decrease in active histone methylation marks during aging that is not explained by changes in the expression of the methyltransferase and demethylase enzymes. Further, these data suggest that the global decrease in H3K4me3 and H3K36me3 observed in aging fly heads likely reflects, at least in part, decreases in levels of these active histone marks in the highly abundant photoreceptor neurons.

### Age-associated decreases in H3K4me3 and H3K36me3 do not correlate with changes in differential gene expression

Because H3K4me3 and H3K36me3 levels correlate with gene expression (transcript levels by RNA-seq) in young photoreceptors ^15^, we next asked if the genome-wide decreases in the active histone methyl marks correlated with changes in gene expression. To do this, we compared our data with the genes that we previously identified as being differentially expressed between day 10 and day 40 in photoreceptors ^25^. In that analysis, we had found that 13% of the 8680 expressed genes in photoreceptors were differentially expressed with age (adjusted p-value < 0.05, |FC| < 1.5), with 570 and 599 genes being down- or up-regulated, respectively between day 40 and day 10 (Fig. 2A). When we plotted H3-normalized H3K4me3 or H3K36me3 levels over the TSS (H3K4me3) or gene body (H3K36me3) for age up- or down-regulated genes relative to genes that were not differentially expressed (no change), we found that actively expressed genes showed age-associated decreases in H3K4me3 and H3K36me3 regardless of their differential gene expression status during aging (Fig. 2B). Moreover, when we examined chromatin accessibility changes obtained using ATAC-seq for differentially expressed genes, we found that there were modest decreases in chromatin accessibility around the TSS during aging at all genes, irrespective of their changes in gene expression (Fig. 2B, right panel). Supporting this decrease in overall levels of chromatin accessibility, we found that H3 levels slightly increased around the TSS when H3 ChIP-seq signal was normalized to input (Fig. 2C). We note that this increase in H3 is not sufficient to explain the decrease in H3K4me3 or H3K36me3 observed in aging, which are observed further downstream of the TSS.

These data argue that the decrease in H3K4me3 and H3K36me3 levels in aging photoreceptors does not directly contribute to changes in differential gene expression. However, we cannot eliminate the possibility that there are global decreases in transcription as photoreceptors age. The modest decrease in chromatin accessibility observed at the TSS at all expressed genes during aging indicates that there might indeed be some global decrease in active transcription with age. However, our current NIE-coupled RNA-seq approach is not compatible with spike-in controls due to the small amount of RNA used for generating libraries (10-30 ng total nuclear RNA), so we are unable to determine if there are global decreases in transcription in aging photoreceptors using these data. Despite these caveats, our data suggest that the deposition and/or maintenance of these active histone methyl marks becomes defective with increasing age in photoreceptors, raising the question as to whether this has any impact on other chromatin-associated processes.

### Elucidating the transcriptional response to decreased Set2/H3K36me3 levels in photoreceptors

To directly test the impact of decreased levels of a histone mark on the transcriptome of photoreceptor neurons, we sought to perform nuclear RNA-seq in photoreceptors with depleted H3K36me3. We co-expressed GFP^KASH^ with a shRNA that targets the Set2 transcript for degradation, enabling us to perform RNA-seq in photoreceptors with decreased H3K36me3. We focused on SET domain containing 2 (*Set2-FBgn0030486*) because it is the sole histone methyltransferase in charge of depositing the third methyl group on H3K36 in *Drosophila* ^20^. In contrast, H3K4me3 can be deposited by three different methyltransferases, SET domain containing 1 (*Set1-FBgn0040022*), trithorax-related (*trr-FBgn0023518*), and trithorax (*trx-FBgn0003862*) ^19^.

We first tested efficiency of the shSet2 in larvae using the ubiquitous *tub-Gal4* driver, and observed significant decreases in both Set2 transcript levels using qPCR (Fig. 3A) and H3K36me3 levels relative to shControl, mCherry RNAi, as measured by western blot (Fig. 3B). To knock down Set2 specifically in photoreceptors, we next expressed shSet2 under the photoreceptor-specific *Rh1-Gal4* driver ^37^. The *Rh1-Gal4* driver becomes active near the end of pupal development, allowing us to avoid disrupting eye development ^37^. To test efficacy of the shRNA knockdown in photoreceptors, we monitored levels of photoreceptor-specific *Rh1-luciferase* in flies expressing shRNA against either luciferase or mCherry control. These data show that shRNA expressed under *Rh1-Gal4* become effective in adults by 2 days post-eclosion and remains effective throughout aging (Fig. 3C).

**Figure 3.**
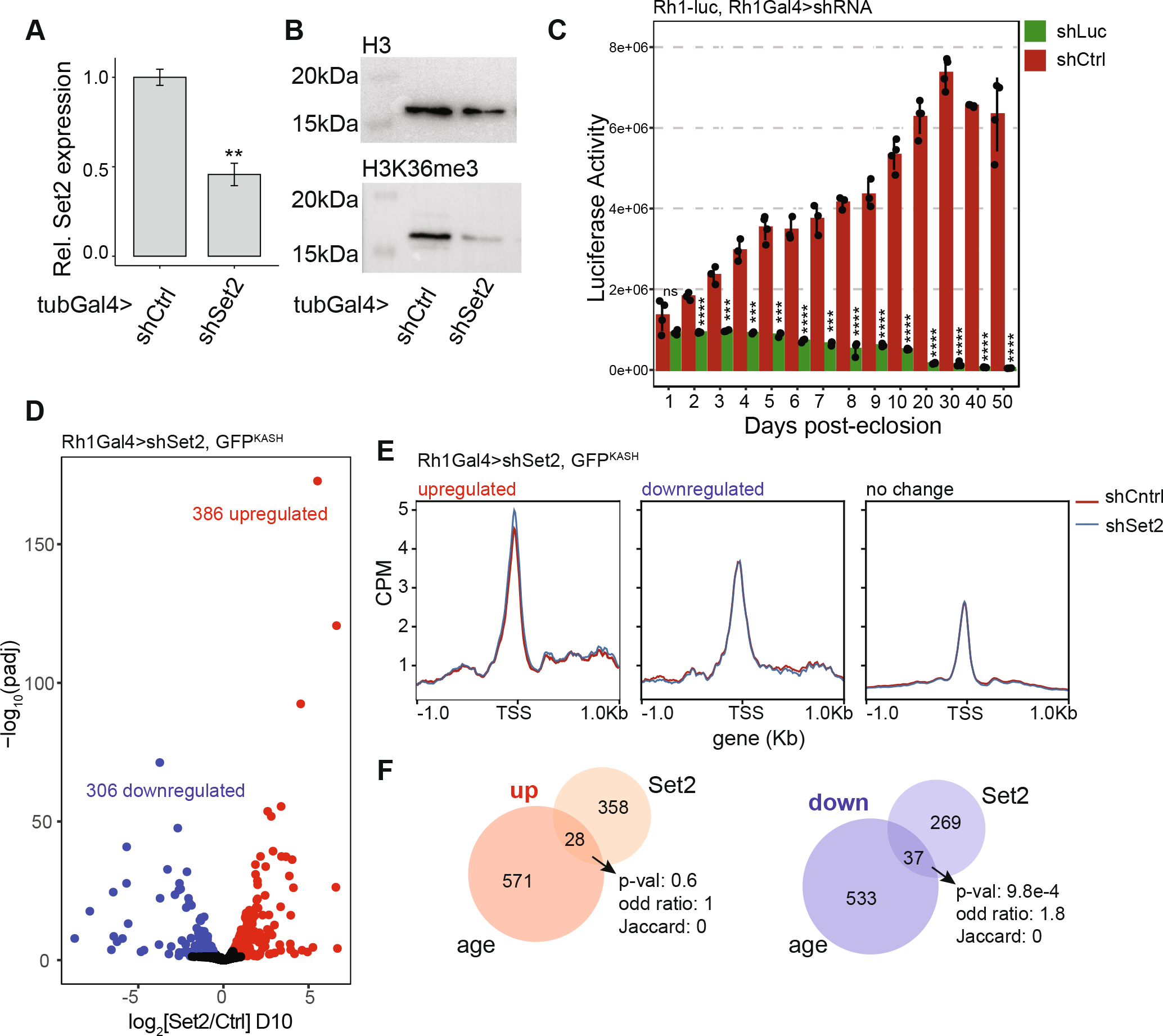
Photoreceptor transcriptome response to Set2 knockdown. **A**. Bar plot showing qPCR analysis of Set2 transcript levels in larvae expressing a shRNA against Set2 (shSet2) or mCherry (shCtrl) under tubGal4 control (mean ± sdev; n = 3). Set2 transcript levels were normalized to the geometric mean of two reference genes (**, p < 0.05, *t*-test). **B**. Western blot showing H3K36me3 and H3 levels in shSet2 and shCtrl larvae, as in panel A. **C**. Bar plot showing luciferase activity in heads from Rh1-luciferase expressing flies that also express a shRNA against luciferase (shLuciferase) or shCtrl under Rh1Gal4 control at the indicated ages in days post-eclosion (mean ± sdev with dots representing individual biological replicates; ns, not significant, ***, p < 0.001, ****, p < 0.0001, *t*-test, n=3). **D**. Volcano plot illustrating genes that were identified as differentially expressed in photoreceptors expressing shSet2 relative to shCtrl. **E**. Gene metaplots of ATAC-seq signal in shSet2 and shCtrl photoreceptors in which genes are classified based on their differential expression status in the Set2 knockdown, as described in panel D. **F**. Venn diagram representing the overlap of genes that were differentially expressed (up – or down-regulated) in either aging (D40 vs D10) or upon loss of Set2 (shSet2 vs shCtrl). Overlap significance was obtained using a hypergeometric test with Jaccard index and Odds ratio are measurements representing similarity.

We next asked how decreased Set2 levels affected gene expression in photoreceptors, to test if decreased H3K36me3 had any contribution to changes in age-associated gene expression. We performed NIE-coupled RNA-seq of photoreceptors expressing shSet2 versus shControl in 10-day-old flies, when knockdown is efficient (Fig. 3C) but flies are still relatively young. Using DESeq2, we performed differential gene expression analysis and found that that 8.8% of actively transcribed genes were differentially expressed in shSet2 relative to shControl, corresponding to 386 and 306 up- and down-regulated genes, respectively (adjusted p-value < 0.05, |FC| < 1.5) (Fig. 3D, Supp. Table 1), suggesting that the photoreceptor transcriptome responds to differential levels of H3K36me3.

To test if these Set2-dependent changes in gene expression were regulated at the chromatin level, we next performed ATAC-seq in Set2-depleted photoreceptors under the same conditions as RNA-seq. Set2/H3K36me3 regulates chromatin accessibility over gene bodies in *Saccharomyces cerevisiae* by recruiting histone deacetyl transferase complexes (HDAC) ^38–40^. Although in *Drosophila* loss of Set2 leads to increased histone acetylation ^20^, ATAC-seq analysis of Set2-null or methylation-deficient H3K36R *Drosophila* larvae showed no changes in accessibility over gene bodies ^41^. Similarly, when we analyzed our ATAC-seq signal over gene bodies, we did not observe any changes in shSet2 relative to shCtrl (Supp. Fig. 3), corroborating these previous observations. Thus, we instead examined the TSS-centered ATAC-seq signal, which positively correlates with transcript levels ^15^, to compare changes in accessibility around the TSS for genes that were differentially expressed in shSet2 flies relative to shControl. At the TSS, we found that genes that were up-regulated at the transcript level, also showed higher accessibility (Fig. 3E). Thus, our data shows that loss of Set2/H3K36me3 leads to differential changes in active transcription. However, when we compared the Set2-dependent transcriptome with age-associated genes, we observed very little overlap (Fig. 3F), consistent with the lack of correlation between differential gene expression and the H3K36me3 decrease in aging photoreceptors. Because H3K36me3 is deposited after the passage of the elongating RNA polymerase II, it is unlikely that Set2 depletion directly contributes to changes in chromatin structure at the TSS, but rather plays an indirect role in transcriptional initiation regulation. Further, we conclude that the age-associated decrease in H3K36me3 in photoreceptors is not a major contributor to changes in differential gene expression or overall chromatin accessibility during aging.

### Set2/H3K36me3 contributes to the aging splicing landscape of photoreceptors

We next wondered if the decrease in H3K36me3 in aging photoreceptors might contribute to other chromatin-associated processes. Deposition of H3K36me3 has been shown to impact splicing by regulating exon selection ^10,42^, and we have previously shown that aging photoreceptors undergo substantial changes in alternative splicing that contribute to decreased visual behavior ^26^. Thus, we asked how loss of H3K36me3 in photoreceptors would impact splicing. Although our NIE-coupled RNA-seq approach includes intronic reads because it is a mixture of nuclear spliced and nascent transcripts, the majority of reads obtained are from spliced transcripts enabling us to examine splicing events in these data ^26^. We performed splicing analysis using rMATS ^43^, which has been shown to out-perform other splicing analysis packages ^44^ including the JunctionSeq software that we previously used to analyze splicing in aging photoreceptors ^26^. Notably, rMATS has the ability to detect different type of splicing outcomes, such as skipped exons and retained introns. Using rMATS, we identified 385 differential splicing events upon loss of Set2, including 146 skipped exon events (FDR<0.05) (Fig. 4A, Supp. Table 2). When we analyzed splicing in young versus old photoreceptors (day 10 versus day 40), we identified 390 splicing events including 161 skipped exons (FDR<0.05) (Fig. 4A, Supp. Table 3). Strikingly, when we compared the gene targets of these differential splicing events that occurred upon loss of Set2 or during aging, we observed a substantial overlap of 123 genes (Fig. 4B). In particular, the gene targets of skipped exon events exhibit a significant overlap with 92 common gene targets (adjusted p-value: 1.50E-56) (Fig. 4C). Skipped exon events are indicative of changes in exon selection, which has been shown to be regulated by Set2 ^10,42^, and can result in proteins with alternative functions. We note that many of the gene targets of these differentially regulated skipped exon events have key roles in photoreceptor function such as the calcium channel *trpl* or the G-protein complex subunit *Ggamma30A* that are involved in phototransduction (Fig. 4A).

**Figure 4.**
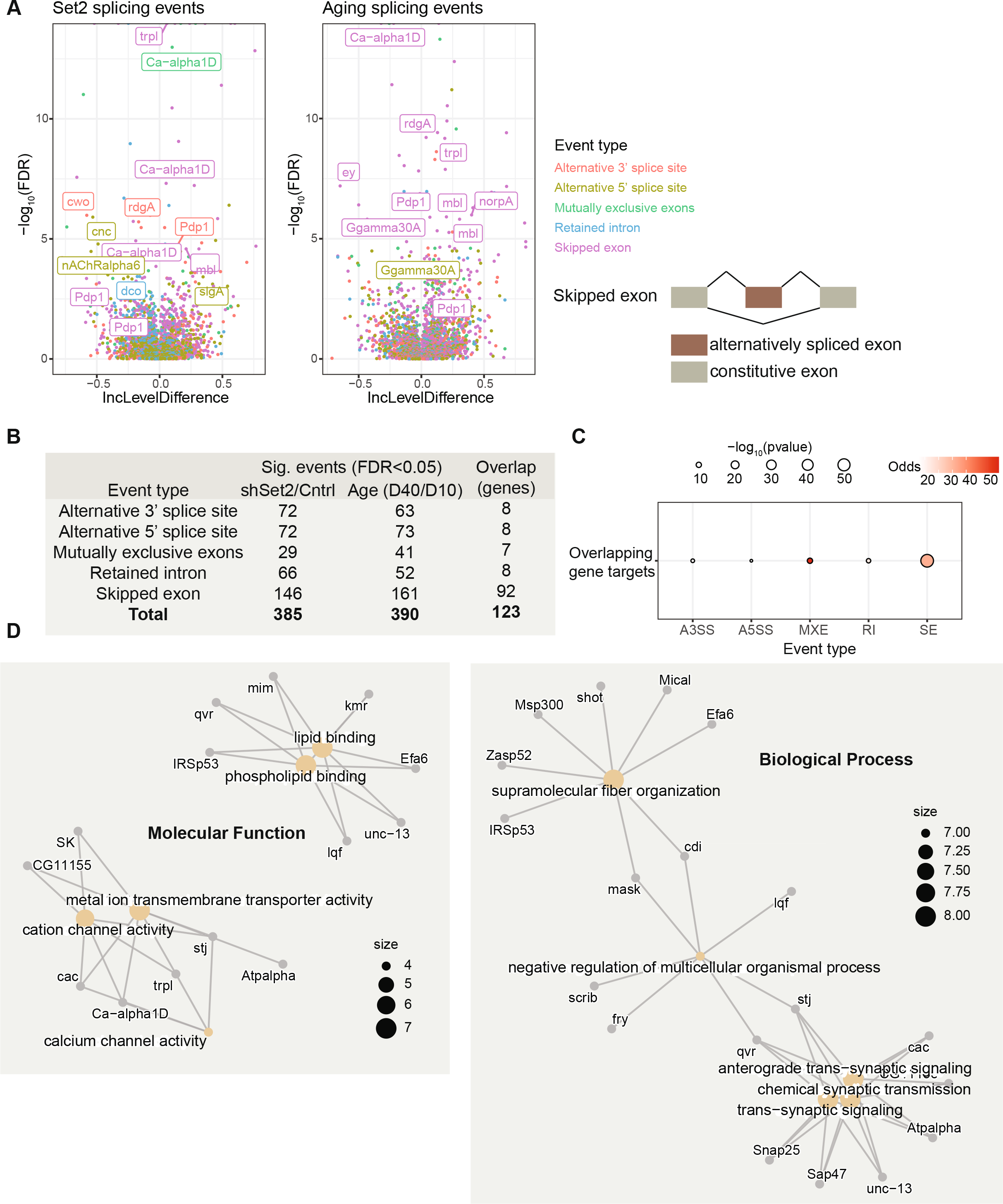
Set2 knockdown leads to differential splicing outcomes that resemble those observed in aging photoreceptors. **A**. Volcano plot representing the differential splicing events identified in shSet2 versus shCntrl at day 10 (left) or aging (day 10 versus day 40) photoreceptors using rMATS. Splicing events are colored by event type, and genes important for photoreceptor function are highlighted. A schematic illustrating skipped exon events is shown to the right. **B**. Table showing the total number of statistically significant events in each category identified using rMATS in aging (D40 vs D10) or upon loss of Set2 (shSet vs shCtrl). Overlapping gene targets for each event are shown in the right column. **C**. Dot plot representing the overlap analysis of splicing events between aging and Set2-deficient photoreceptors. Statistical significance is obtained performing a hypergeometric test. A higher Odds ratio represents similarity between datasets, with values below 1 indicating no association **D**. Gene Concept Network (cnet) plots illustrating individual genes driving the statistically significant enrichment of GeneOntology (GO) Biological Process and Molecular Function terms identified for the shared splicing events for Set2 and aging.

To test if any biological process was enriched among these Set2- and age-regulated splicing events, we performed gene ontology (GO) term analysis. We found that gene targets of the overlapping splicing events were significantly enriched for biological processes and molecular functions important for photoreceptor function (Fig. 4D). These enriched GO terms included calcium channel activity, which is critical for phototransduction in photoreceptors, and synaptic signaling. Notably, the overlapping splicing events also impacted several genes that regulate gene expression processes in photoreceptors including the circadian regulator PAR domain protein 1 (*Pdp1-FBgn0016694)* and the splicing/circRNA binder muscleblind (*mbl*). We previously observed changes in the activity of the circadian clock transcription factors Clock and Cycle in aging photoreceptors ^25^, and these data raise the intriguing possibility that alterations in histone methylation could impact the clock via splicing of circadian regulators. Moreover, our data suggest that loss of Set2 leads to changes in splicing that could have broad and indirect effects on gene expression because they affect proteins with key roles in regulating transcription and splicing. We propose that the decrease in H3K36me3 in aging photoreceptors contributes to changes in alternative splicing outcomes, with potential functional consequences for the function and integrity of photoreceptor neurons.

## DISCUSSION

Here, we show *Drosophila* photoreceptors undergo an age-associated decrease in two histone methylation marks associated with active transcription: H3K4me3 and H3K36me3. These age-dependent decreases in histone methylation were observed at all actively expressed genes and did not correlate with differential gene expression or chromatin accessibility. Although we cannot exclude the possibility that there are global decreases in transcription during aging, our analysis of photoreceptors with Set2 knockdown suggests that H3K36me3 does not directly contribute to age-dependent changes in gene expression changes or chromatin accessibility. Instead, we observe a significant overlap in splicing events between Set2 knockdown and aging photoreceptors, indicating that the decrease in H3K36me3 observed in aging photoreceptors contributes to age-dependent changes in alternative splicing. These data further suggest that the decrease in these two histone methylation marks in aging photoreceptors could have complicated and indirect effects caused by the interplay between these marks, and by Set2-dependent alterations in splicing of gene expression regulators.

Our analysis of aging photoreceptors largely resembles the observations previously reported for fly heads using ChIP-CHIP ^22^, and is similar to reports for aging whole organism *Caenorhabditis elegans* ^29^ and *S. cerevisiae* ^45^ using ChIP-seq. Some studies examining muscle tissue in flies have reported increases in the heterochromatic H3K27me3 mark with age ^23^, but no changes in H3K27me3 were observed in aging heads ^22^. However, age-associated alterations in the distribution of other heterochromatic methyl marks such as H3K9me3 have been reported in heads ^22^. In this study, we focused on only two histone methylation marks associated with active transcription, H3K4me3 and H3K36me3. Thus, our work raises an important question as to whether the decrease in histone methylation is limited to marks associated with active transcription, potentially reflecting decreased recruitment or transcription by RNA polymerase II ^22^. Alternatively, the decrease in histone methylation might reflect changes in the activity or recruitment of the enzymes that deposit and remove these marks, although we did not observe changes in their expression at the transcript or protein level. The modest decrease in chromatin accessibility around the TSS in aging photoreceptors suggests that there might be an overall decrease in transcription as these neurons age, but it remains to be determined whether the observed decrease in histone methylation is a cause or consequence of this event.

Our observation that the overlapping splicing events common to Set2 knockdown and aging photoreceptors include many genes associated with phototransduction, and neuronal and photoreceptor function suggests that decreased H3K36me3 levels in aging photoreceptors might have broader long-term effects on photoreceptor homeostasis than initially suggested by the limited effect of Set2 knockdown on gene expression. Supporting this, we previously showed that proper splicing is critical for maintaining visual behavior and function in the aging *Drosophila* eye ^26^. Although we focus on skipped exons i.e. alternative splicing events in this study, we previously showed that aging photoreceptors accumulate circRNAs ^4^, which are generated from back-splicing events ^46^. H3K36me3 has been implicated as a negative regulator of circRNA levels ^47^, and one of the mRNAs exhibiting skipped exons under Set2-knockdown and aging encodes muscleblind (mbl), which sequesters circRNAs, and has eye- and brain-specific functions ^48^. These data suggest one mechanism through which H3K36me3 decreases could contribute to more widespread alterations in the aging transcriptome, by impacting biogenesis of circRNAs, as well as the activity of circRNA binding proteins. Another overlapping splicing target between aging and Set2 knockdown was Pdp1, a component of the circadian clock that maintains the ~24h periodicity of many biological processes ^49^. Activity of the circadian clock transcription factors alters with age in photoreceptors, and disrupting the circadian clock in the fly or mouse eye results in progressive retinal degeneration ^25,50^. Since we only examined one time point during the day in our RNA-seq studies, our data are insufficient to reveal any effect of Set2 depletion on circadian-regulated gene expression. However, alternative splicing of several core clock mRNAs affects circadian behavior ^51–56^, suggesting that H3K36me3-dependent splicing events could impact the circadian clock. Alternatively, the decrease in H3K4me3 in aging photoreceptors could play a more important role in circadian-dependent gene expression. The mammalian H3K4me3 methyltransferase MLL1 (trx in flies) is required for expression of circadian clock output genes in mouse embryonic fibroblasts, and directly interacts with the transcription factors CLOCK-BMAL1 to deposit H3K4me3 ^57^. Moreover, in flies, loss of the histone H3K36me3 demethylases Kdm2 and Jmjd5, or the H3K4me3 demethylase Lid (human JARID1A), alters circadian behaviors ^58–61^. JARID1A and JMJD5 are also necessary for circadian rhythms in mammals ^62,63^. Comprehensive examination of the transcriptome over multiple time points during the day during aging will be necessary to determine how alterations in these histone methyl marks might impact circadian regulated gene expression.

Overall, our studies here show that aging photoreceptors experience a widespread decrease in chromatin modifications associated with active transcription. These findings are not unique to *Drosophila*, since analysis of aging tissues for *C. elegans*, mice and human have shown similar widespread changes in histone modifications together with changes in histone variants (Pal and Tyler 2016; Kane and Sinclair 2019). While we found no strong correlation between these changes in histone modifications and differential gene expression, our analysis suggests that H3K36me3 has an important role in regulating splicing outcomes in aging photoreceptors. Since differential splicing has been recently linked to aging and neurodegenerative disorders ^42,64,65^, our studies suggest that H3K36me3 and Set2 could be interesting targets for modulation in the context of retinal and neurodegenerative disease.

## MATERIALS AND METHODS

### Drosophila

*Drosophila* stocks were raised and NIE and ChIP-seq performed as previously described ^15,66^. The following genotypes were used: Rh1-Gal4>UAS-GFP^KASH^ (*w^1118^;; P{w^+mC^=[UAS-GFP-Msp300KASH}attP2, P{ry^+t7.2^=rh1-GAL4}3, ry^506^*); Set2 RNAi (BDSC #42511; *y^1^ v^1^; P{y^+t7.7^ v^+t1.8^=TRiP.HMJ02076}attP40*); mCherry RNAi (*yv; P{w^v+^=[UAS-mCherry-RNAi]}attP2*) ^26^; Rh1-luciferase (*P{w^+mC^=Rh1-Ffluc}3*) ^26^.

### Quantitative PCR

qPCR was performed as previously described ^15^ with the following primers:*eiF1α (5’*- GCTGGGCAACGGTCGTCTGGAGGC-3’ and 5’- CGTCTTCAGGTTCCTGGCCTCGTCCGG-3’); *Rpl32* (5’- GCTAAGCTGTCGCACAAATG-3’ and 5’- CGTTGTGCACCAGGAACTT –3’); *Set2* (5’- GGTCAAGAGTTCCCAGAGTCC-3’ and 5’- CTCATCGCTGGTGATCCGAC-3’).

### Western blot and luciferase assays

Histones were extracted using a modified acid extraction protocol. Briefly, five third instar larvae were homogenized in 200 μL of Extraction buffer (1X PBS, 0.5% Triton X-100, 2 mM PMSF) using a motorized pestle motor, and incubated for 10 minutes at 4°C with constant rotation. Samples were centrifuged at 6,500 *x g* for 10 min at 4°C and the pellet was resuspended in 100 μL of 0.4N HCl and incubated on ice for 5 minutes, followed by centrifugation at 6,500 *x g* for 10 min at 4°C. The supernatant was transferred to a fresh tube and neutralized with NaOH, then analyzed by SDS-PAGE and western blotting using H3 (Abcam; ab1791) and H3K36me3 antibodies (Abcam; ab9050). Luciferase assays were performed as previously described ^26^. Full western blots corresponding to cropped panels are shown in supp Figure 4.

### NIE and ChIP-seq and bioinformatics analysis

NIE and ChIP-seq was performed as previously described ^15^. Detailed ChIP-seq instructions are available at: https://www.protocols.io/view/in-vivo-tissue-specific-chromatin-profiling-in-dro-kxygxp1d4l8j/v1.ChIP-seq and RNA-seq analysis was performed as previously described ^15,25,67^. RNA-seq significance thresholds were based on absolute fold change ≥ 1.5, and adjusted p-value ≤ 0.05. Splicing analysis was performed using rMATS (v4.1.2) ^43^. Heatmaps and gene metaplots were generated using deepTools package plotProfile and plotHeatmap ^68^.

## ACKNOWLEDGEMENTS

We thank the Weake lab for their comments on the manuscript. The authors acknowledge the use of the facilities of the Bindley Bioscience Center, a core facility of the NIH-funded Indiana Clinical and Translational Sciences Institute.

## AUTHOR CONTRIBUTIONS

J.J-L. performed all the Next Generation Sequencing experiments and bioinformatic analysis. K.M.B. performed the quantitative PCR. A.C.H. performed the western blot. J.J-L. and V.M.W wrote and edited the manuscript, with input from other authors.

## DATA AVAILABILITY STATEMENT

Photoreceptor ChIP-seq data for aging H3, H3K4me3, and H3K36me3 is available from the Gene Expression Omnibus (GEO) repository under series accession number GSE202050. RNA-seq and ATAC-seq data for Set2 knockdown photoreceptors versus control are available under GSE202053 and GSE174491. Previously published RNA-seq and ATAC-seq expression data are accessible under: GSE169328 (D10 ATAC-seq and RNA-seq), GSE184069 (D40 ATAC-seq) and GSE174515 (D40 RNA-seq). Flies carrying UAS-GFP^KASH^, as well as additional flies with different KASH-tagged epitopes are available at Bloomington Drosophila Stock Center. Step-by-step protocols are available at dx.doi.org/10.17504/protocols.io.buiqnudw.

