## Supplemental Figures for "Establishing the contribution of active histone methylation marks to the aging transcriptional landscape of Drosophila photoreceptors"

**SUPPLEMENTAL DATA:**

**Supplemental Figure 1. A.** Experimental diagram of chromatin immunoprecipitation coupled with incorporation of exogenous *Arabidopsis* chromatin for spike-in normalization. **B.** PCA of spike-in and H3-normalized H3K4me3 and H3K36me3 samples during aging. **C.** Gene metaplots showing individual replicates used to compute the average gene metaplot in Figure 1D. **D.** Gene metaplots showing spike-in and H3-normalized H3K4me3 and H3K36me3 samples during aging.

**Supplemental Figure 2.** Boxplots of expression values (TPM, transcripts per million relative to day 10, set at one) for H3K4me3 and H3K36me3 writers and erasers. Differential expression analysis was used to determine statistically significant changes in gene expression between D10 and D40 samples. ns., not significant, * adj.p-val<0.05, ** adj.p-val<0.005.

**Supplemental Figure 3. A.** Gene metaplot showing ATAC-seq signal (CPM) over gene bodies in photoreceptors expressing shSet2 versus shControl at day 10. **B.** MA plot showing significant changes in chromatin accessibility determined by ATAC-seq at peaks in photoreceptors expressing shSet2 versus shControl at day 10.

**Supplemental Figure 4.** Full panels for western blot showing H3K36me3 and H3 levels in larvae expressing a shRNA against Set2 (shSet2) or mCherry (shCtrl) under tubGal4 control as shown in Figure 3B.

**Supplemental Table 1:** Differentially expressed genes in shSet2 photoreceptors versus shControl.

**Supplemental Table 2:** Differential splicing events in shSet2 photoreceptors versus shControl.

**Supplemental Table 3:** Differential splicing events in day 10 versus day 40 photoreceptors.

Jauregui-Lozano_suppFig1

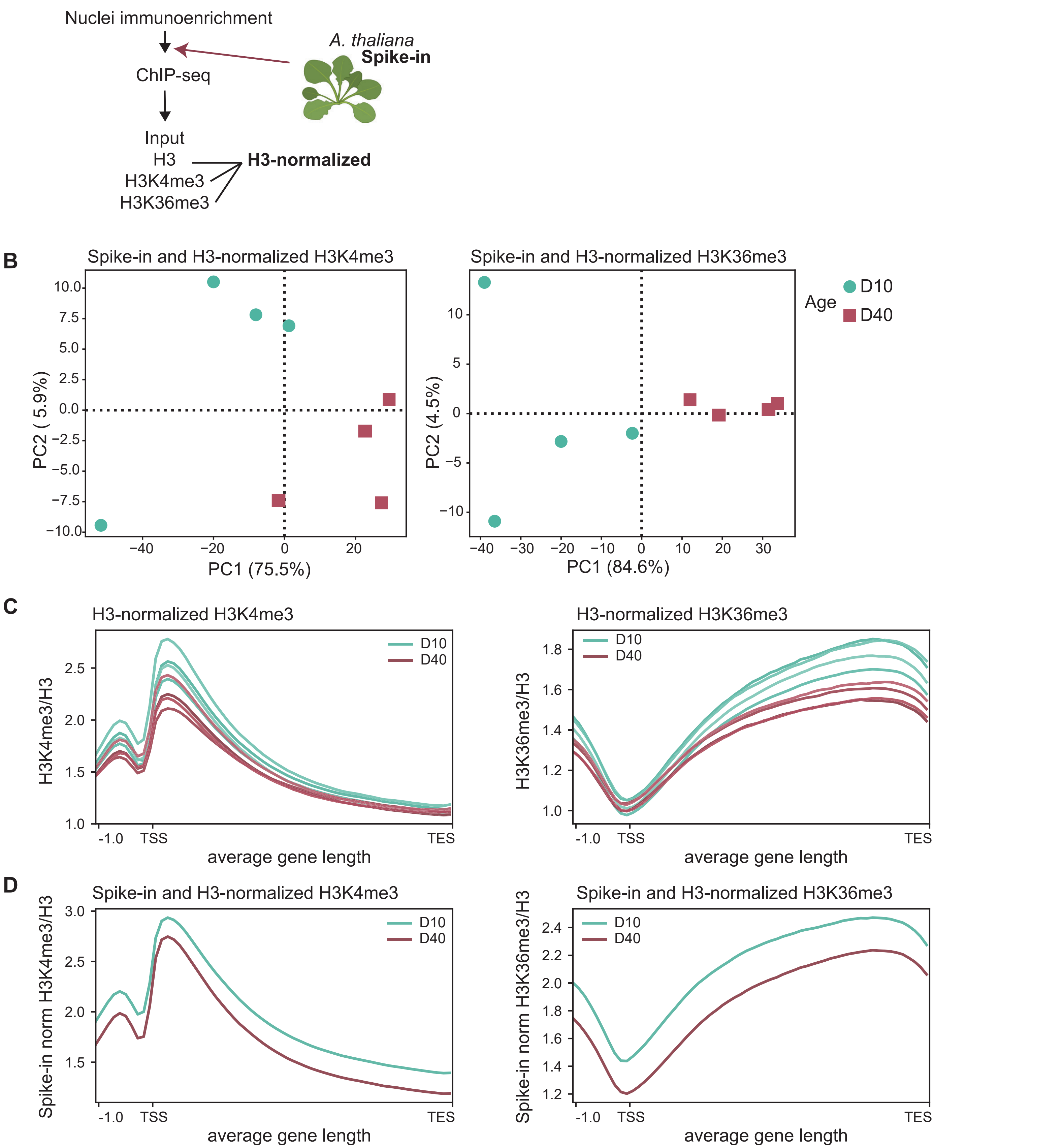

Jauregui-Lozano_suppFig2

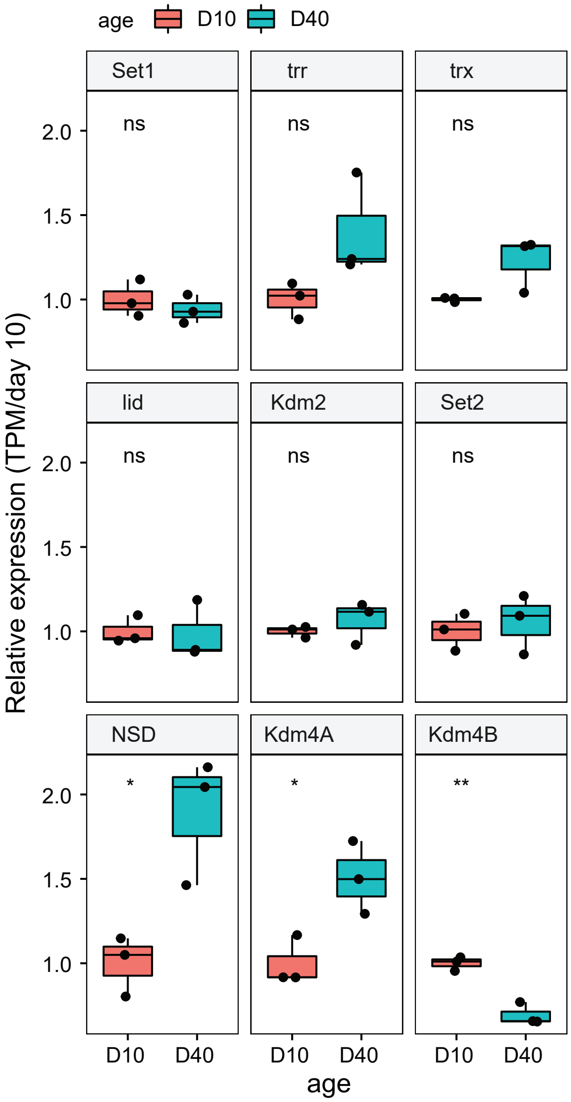

Jauregui-Lozano_suppFig3

A

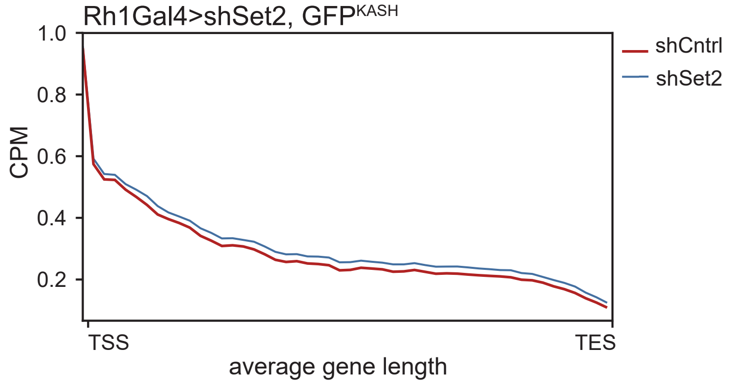


B

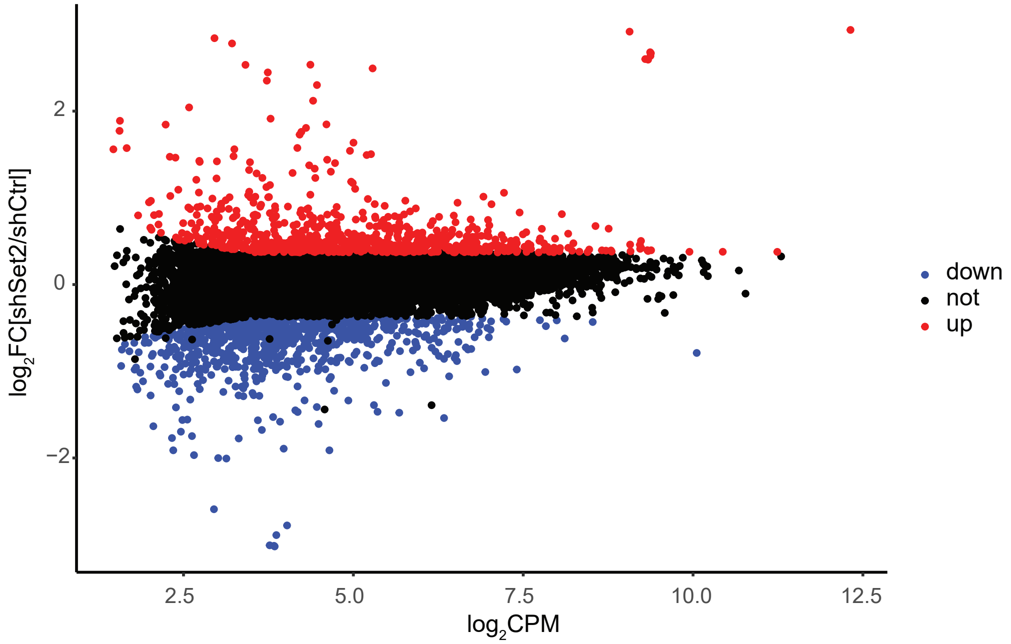

Jauregui-Lozano_suppFig4

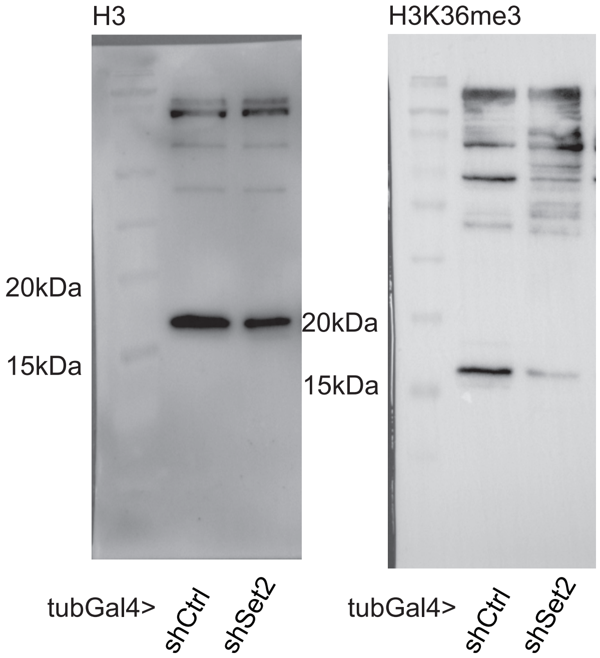
